# PhredEM: A Phred-Score-Informed Genotype-Calling Approach for Next-Generation Sequencing Studies

**DOI:** 10.1101/046136

**Authors:** Peizhou Liao, Glen A. Satten, Yi-juan Hu

**Affiliations:** Department of Biostatistics and Bioinformatics, Emory University, Atlanta, Georgia; Centers for Disease Control and Prevention, Atlanta, Georgia

**Keywords:** common variant, EM algorithm, low depth, rare variant, read data

## Abstract

A fundamental challenge in analyzing next-generation sequencing data is to determine an individual’s genotype correctly as the accuracy of the inferred genotype is essential to downstream analyses. Some genotype callers, such as GATK and SAMtools, directly calculate the base-calling error rates from *phred* scores or recalibrated base quality scores. Others, such as SeqEM, estimate error rates from the read data without using any quality scores. It is also a common quality control procedure to filter out reads with low *phred* scores. However, choosing an appropriate *phred* score threshold is problematic as a too-high threshold may lose data while a too-low threshold may introduce errors. We propose a new likelihood-based genotype-calling approach that exploits all reads and estimates the per-base error rates by incorporating *phred* scores through a logistic regression model. The algorithm, which we call PhredEM, uses the Expectation-Maximization (EM) algorithm to obtain consistent estimates of genotype frequencies and logistic regression parameters. We also develop a simple, computationally efficient screening algorithm to identify loci that are estimated to be monomorphic, so that only loci estimated to be non-monomorphic require application of the EM algorithm. We evaluate the performance of PhredEM using both simulated data and real sequencing data from the UK10K project. The results demonstrate that PhredEM is an improved, robust and widely applicable genotype-calling approach for next-generation sequencing studies. The relevant software is freely available.

## INTRODUCTION

The recent advancement of next-generation sequencing (NGS) technologies and the rapid reduction of sequencing costs have led to extensive use of sequencing data in disease association studies and population genetic studies [Ng et al., 2010; The 1000 Genomes Project Consortium, 2010]. However, it is still difficult and costly to perform whole-genome sequencing (WGS) with high depth in large cohorts [Sims et al., 2014]. Instead, many studies have adopted whole-exome sequencing (WES) [The 1000 Genomes Project Consortium, 2012; Muddyman et al., 2013]. Despite the high average depth that is typically attainable in WES studies, some regions within a gene may still have much lower depth than the average due to the inefficiency of exome capture technologies [Do et al., 2012]. Other studies have kept the design of WGS but chosen low or moderate depths. For example, the UK10K project (www.uk10k.org) sequenced the two population cohorts genome wide at depth of ~6x. Although sequencing costs are declining, we anticipate that many NGS studies will continue to employ WES or WGS with low or medium depth for some time to come.

A fundamental challenge in analyzing NGS data is to determine an individual’s genotype correctly, as the accuracy of the inferred genotype is essential to downstream analyses. It is difficult to call genotypes for two reasons. First, NGS data can suffer from errors introduced in the base-calling process. The base-calling error rate ranges from a few tenths of a percent to several percent [Nielsen et al., 2011]. It can vary from base to base as a result of machine cycle and sequence context [Kircher et al., 2009]. It also varies dramatically across different sequencing platforms. For instance, the Illumina MiSeq platform has an error rate of ~0.8% [Quail et al., 2012] whereas the Roche 454 System has ~0.1% [Liu et al., 2012]. Second, the quality of called genotypes depends heavily on the read depth. Genotypes covered by many reads can typically be called reliably. However, when a locus is covered by only a few reads, genotype calling is challenging because minor allele reads are indistinguishable from sequencing errors.

*Phred* scores are widely accepted to characterize error rates in the base-calling process.

All major sequencing platforms assign each called base of a raw sequence a *phred* score, which measures the probability that the base is called incorrectly [Ewing et al., 1998; Ewing and Green, 1998]. *Phred* scores are determined using various predictors of possible errors such as peak spacing, uncalled/called peak ratio and peak resolution. Nominally, the *phred* score is defined as

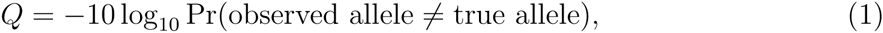

so that, for example, *Q* = 30 nominally corresponds to a 0.1% error rate. Despite their wide use, *phred* scores may not accurately reflect the true error rates in base calling because they fail to account for some important factors. For instance, the specific error pattern inherent in each nucleotide base (i.e., A, C, T and G) is not considered in *phred* scores [Li et al., 2004]. Additionally, *phred* scores do not account for the position of the base within a read [DePristo et al., 2011]. Since *phred* scores might be inaccurate representation of true base-calling error rates, many methods have been developed to recalibrate base quality scores [DePristo et al., 2011; Li et al., 2009b]. However, although recalibrated scores could be more accurate than *phred* scores, the recalibration process may be too computationally intensive to be of broad practical use [Nielsen et al., 2011].

A genotype-calling method typically uses a probabilistic framework, combining base-calling error rates and a prior distribution of genotype frequencies to provide a posterior probability for each genotype [Mckenna et al., 2010; Li et al., 2009a; Martin et al., 2010]. Because the error rate is critical in probabilistic genotype-calling algorithms, it is crucial that it be correctly specified, especially when sequencing depth is low to moderate. Some methods such as GATK use error rates that are calculated directly from *phred* scores by applying equation (1) if recalibration step is skipped. In contrast, SAMtools obtains an error rate from the minimum of the *phred* score and the mapping score [Li, 2011]. In addition, bases with low *phred* scores (e.g., *Q* < 20 or 30) are typically filtered out as part of quality control (QC) procedures. However, there are some concerns in choosing a threshold for *phred* scores. High thresholds may result in loss of useful information by eliminating bases that are correctly called. Low thresholds leave a large number of erroneously called bases in the data, leading to false-positive variant calls.

Instead of relying on *phred* scores, Martin et al. [2010] proposed SeqEM, a genotype-calling algorithm that estimates the error rate using the read data itself. However, the fundamental assumption of SeqEM that at each locus a uniform error rate exists for all bases across the sample is generally not true, given the considerable variability in error rates implied by the variability in *phred* scores. Also, as SeqEM ignores *phred* scores entirely, the valuable information about errors encoded in *phred* scores is lost.

In this paper, we propose a new genotype-calling approach which estimates base-calling error rates from the read data while incorporating the information in *phred* scores. We model an error rate as a logistic function of a *phred* score; this logistic regression model is readily integrated into a modification of the SeqEM likelihood which allows for a base-specific error probability. Like SeqEM, our approach also uses the Expectation-Maximization (EM) algorithm [Dempster et al., 1977]. Information from all individuals is used to estimate the unknown genotype frequencies and logistic regression parameters. We compute the posterior probability of each latent genotype based on parameter estimates and use the empirical Bayes approach to assign the most likely genotype to each individual. We show that the logistic model fits real sequencing data well, and that the unknown parameters in our likelihood are consistently estimated. Moreover, to minimize the effort of calling genotypes for loci with no variation, we develop a simple, computationally efficient screening algorithm to identify loci that are estimated to be monomorphic (and therefore do not require parameter estimation using the EM algorithm). Finally, we demonstrate through simulation studies that our approach is more accurate than SeqEM. We illustrate our new approach through an application to real sequencing data from the UK10K project.

## METHODS

We consider one biallelic locus at a time. For the *i*-th individual, let *G*_*i*_ denote the underlying true genotype (coded as the number of minor alleles), *T*_*i*_ denote the total number of alleles that are mapped to the locus, and *R*_*i*_ (*R*_*i*_ ≤ *T*_*i*_) denote the number of mapped alleles that are called to be the minor allele. The *phred* scores are represented by ***Q***_*i*_ = (*Qi*1,…, *Q-iTi*′, where *Q*_*ik*_ is the *phred* score associated with the *k*-th called allele and the prime (′) indicates the transpose of a vector. At each locus, values of *T*_*i*_, *R*_*i*_, and ***Q***_*i*_ can be easily extracted from the pileup files produced by SAMtools. Let *ϵ*_*ik*_ be the true base-calling error rate of the *k*-th allele. We relate *ϵ*_*ik*_ to *Q*_*ik*_ through the logistic regression model

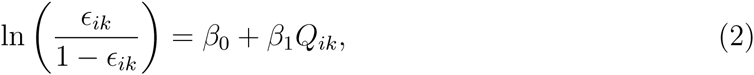

where *β*_0_ and *β*_1_ are unknown regression parameters that are locus specific. Let *θ* = (*β*_0_, *β*_1_)′ and *ϵ*_*ik*_(*θ*) = exp(*β*_0_ + *β*_1_*Q*_*ki*_)/{1 + exp(*β*_0_ + *β*_1_*Q*_*ik*_)}. Equation (2) is motivated by the fact that the *phred* score is a highly informative predictor of the base-calling error, even though (1)does not hold in the exact sense. In the Results section, we demonstrated that the logistic model fits the real sequencing data well.

Without loss of generality, we order the *T*_*i*_ alleles so that the first *R*_*i*_ alleles are called to be the minor allele and the rest the major allele. Assuming that the errors of the *T*_*i*_ alleles are independent of each other, the probability of observing *R*_*i*_ copies of the minor allele out of *T*_*i*_ alleles can be described as a sequence of independent Bernoulli trials. Specifically, given the true genotype *G*_*i*_, the total number of alleles *T*_*i*_, and the *phred* scores ***Q***_*i*_, the probability of observing *R*_*i*_ is written as

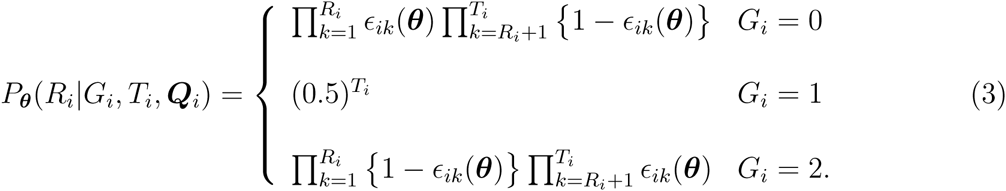

Suppose that the sample consists of n unrelated individuals. Then the likelihood function takes the form

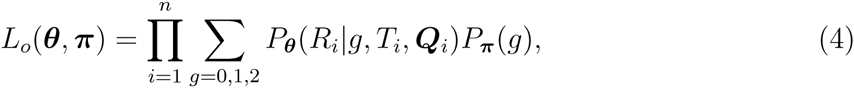

where *P*_*π*_ (*g*) is the genotype frequency characterized by ***π***. Under Hardy-Weinberg Equilibrium (HWE), ***π*** consists of a single parameter *π* for the minor allele frequency (MAF). Then, *P*_*π*_(0) = (1 — *π*)^2^, *P*_*π*_(1) = 2*π*(1 — *π*), and *P*_*π*_(2) = *π*^2^. Under Hardy-Weinberg Disequilibrium (HWD), *π* = (*π*, *f*)′ where *π* and *f* are the MAF and the fixation index *F*_*st*_, respectively. Then, *P*_*π*_(0) = (1 — *f*)(1 — *π*)^2^ + *f* (1 — *π*), *P*_*π*_(1) = 2*π*(1 — *π*)(1 — *f*), and *Pπ* (2) = (1 — *f*)*π*^2^ + *fπ*.

The proposed likelihood is closely related to several existing methods. When *β*_*1*_ = 0, the error rate is independent of the *phred* score, and expression (4) reduces to the likelihood of SeqEM. When *β*_0_ = 0,*β*_1_ = — ln(10)/10, and ***π*** is known, the right hand side of equation (2)becomes — *Q* ln(10)/10. When all error rates are small, which is expected, expression (4) is approximately the likelihood of the Bayesian genotyper implemented in GATK. However, our likelihood fully exploits the read data and the *phred* scores, both of which could improve genotype-calling accuracy. Note that it is not necessary to filter out low-quality alleles, which still provide some information about ***θ**.* Like other multi-sample calling methods, our method also estimates the genotype frequencies and regression parameters by utilizing information across all individuals in the sample.

We obtain estimates of ***θ*** and *π* by maximizing the likelihood (4) via the EM algorithm described in the Appendix. To ensure that increasing *phred* scores correspond to decreasing error rates, we maximize the likelihood subject to the constraint *β*_1_ ≤ 0. Denote the MLEs by ***π̂*** and ***θ̂***. We can estimate the posterior probability distribution of the true genotype *G*_*i*_ from the read count data *T*_*i*_ and *R*_*i*_ and the *phred* scores ***Q***_*i*_ for each study subject according to the formula

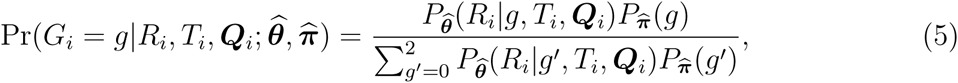

for *g* = 0, 1 and 2. Genotype calls can be made by assigning each individual the genotype with the highest estimated posterior probability. Individuals with no read covering the locus are not assigned any genotype. Because the proposed method incorporates the *phred* scores and uses the EM algorithm, we refer to it as PhredEM.

The majority of loci in the human genome are monomorphic [The International SNP Map Working Group, 2011], and are generally of little interest in downstream analyses. Because it is a waste of time to run PhredEM at such loci, we propose the following simple and computationally efficient algorithm to screen them out without applying PhredEM. We assume HWE holds, because loci that might be called monomorphic must have either zero or extremely low MAFs. We see that formula (5) assigns all mass to *G*_*i*_ = 0 when *π̂* = 0; thus loci with *π̂* = 0 would be called monomorphic if PhredEM was applied to obtain n. We now give a simple way to determine whether *π̂* = 0. Let *pl* (*π*) denote the profile likelihood for *π*, namely,

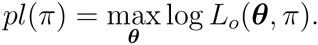

We show in the Appendix that *pl*(*π*) is a concave function of *π*, so that a negative value for the derivative of *pl*(*π*) at *π* = 0 implies *π̂* = 0; in other words, we should screen out loci at which the derivative of *pl*(*π*) at *π* = 0 is negative. At *π* = 0, we can easily evaluate this derivative, because the likelihood *L*_o_(***θ***, *π*) reduces to that of a logistic regression model in which we assign an outcome variable *Y*_*ik*_ = 1 to a minor allele read and *Y*_*ik*_ = 0 to a major allele read and regress *Y*_*ik*_ on *Q*_*ik*_. Since our screening algorithm only involves fitting a standard logistic regression model to solve for ***θ*** and calculating a derivative function, it can significantly reduce the computing time that is needed by the full PhredEM algorithm.

## RESULTS

### SIMULATION STUDIES

We conducted simulation studies to assess the performance of PhredEM relative to SeqEM. We considered a sample size of 1,000 (results based on the sample size of 200 are reported in Supplemental Tables S1 and S2) and three average depths, 6x, 10x, and 30x. For common alleles, we generated loci with a specified allele frequencies, while for rare alleles we generated loci with a fixed number of minor alleles. Based on our analysis of the UK10K data, we generated the depth *T*_*i*_ for the *i*-th individual from the negative-binomial distribution with the given average depth and dispersion parameter 0.35. We then generated the ‘true’ allele corresponding to each read; for heterozygotes the true allele for each read was assigned randomly. Next, for each read, we simulated a *phred* score from the empirical distribution observed in the UK10K data (Figure 1a), calculated the error rate according to (2), and generated a called allele using this error rate. The parameters *β*_0_ and *β*_1_ in (2) were set to be —0.838 and —0.240, respectively, which are median values of their estimates obtained from loci that were determined to be monomorphic by our screening algorithm (so that we could treat all minor allele reads as errors) in analysis of the UK10K data.

**Figure 1:**
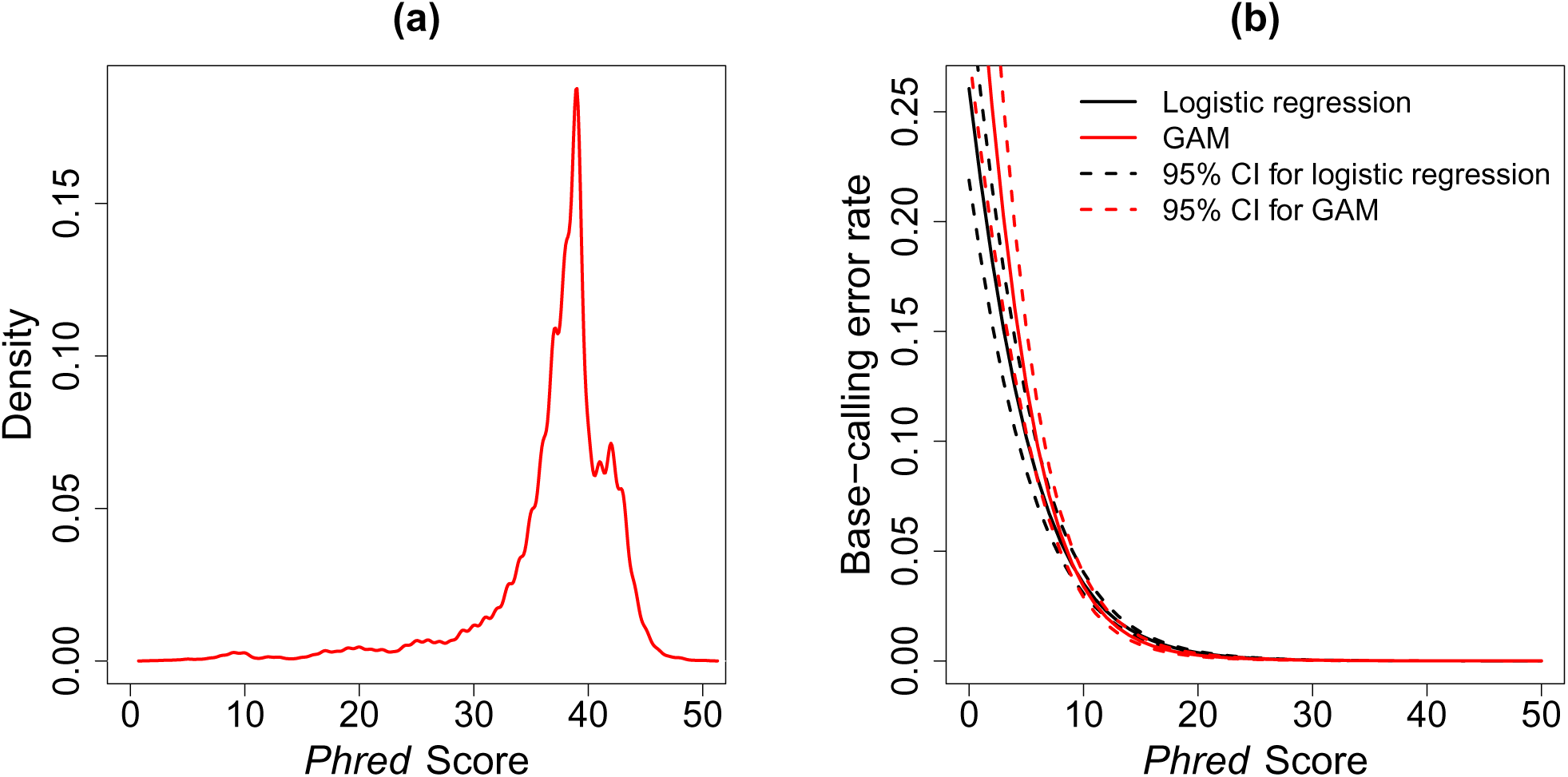
UK10K SCOOP data. (a) Distribution of *phred* scores. (b) Logistic regression model and generalized additive model (GAM) fit to the sequencing data at loci that were identified to be monomorphic.

We first evaluated the performance of PhredEM and SeqEM on rare variants. We generated the true genotypes by fixing the minor allele count (MAC) in each replicate. We considered MACs of 1, 5, 10 and 20, where MAC = 1 corresponds to a singleton. In applying PhredEM and SeqEM, we assumed HWE in both methods, which assumption has a minimal constraint for rare variants because homozygotes of minor alleles are not expected. As shown in Table I, the overall number of mis-called genotypes obtained by PhredEM was less than that by SeqEM in all scenarios. In particular, PhredEM reduced by almost one half the number of mis-called genotypes compared with SeqEM. For instance, when MAC was 10 and depth was 6x, SeqEM mis-called an average of 2.27 genotypes among 1,000 individuals whereas PhredEM mis-called 1.31. As expected, both methods became more accurate as the average read depth increased. Nevertheless, the performance of PhredEM was noticeably better than SeqEM even at a depth as high as 30x. We further examined the mis-called genotypes stratified by the true genotype. In both strata of homozygote (*G* = 0) and heterozygote (*G* = 1), PhredEM mis-called fewer genotypes than SeqEM. At depth of 30x, PhredEM almost detected all rare alleles whereas SeqEM missed one copy of rare allele for every 100 singletons.

**Table I:** Number of mis-called genotypes for rare variants in the simulation studies.

| MAC | Depth | Overall |  | Stratified |  |  |  |
| --- | --- | --- | --- | --- | --- | --- | --- |
| | | P | S | $G = 0$ | | $G = 1$ | |
|  |  |  |  | P | S | P | S |
| 1 | 6x | 0.08 | 0.21 | 0.05 | 0.07 | 0.03 | 0.14 |
|  | 10x | 0.04 | 0.11 | 0.02 | 0.03 | 0.02 | 0.08 |
|  | 30x | 0.01 | 0.02 | 0.01 | 0.01 | 0 | 0.01 |
| 5 | 6x | 0.74 | 1.38 | 0.10 | 0.35 | 0.64 | 1.03 |
|  | 10x | 0.39 | 0.71 | 0.07 | 0.15 | 0.32 | 0.56 |
|  | 30x | 0.05 | 0.10 | 0.01 | 0.02 | 0.04 | 0.08 |
| 10 | 6x | 1.31 | 2.27 | 0.15 | 0.63 | 1.16 | 1.64 |
|  | 10x | 0.67 | 1.20 | 0.10 | 0.31 | 0.57 | 0.89 |
|  | 30x | 0.08 | 0.16 | 0.02 | 0.04 | 0.06 | 0.12 |
| 20 | 6x | 2.31 | 3.62 | 0.20 | 1.00 | 2.11 | 2.62 |
|  | 10x | 1.17 | 1.95 | 0.14 | 0.54 | 1.03 | 1.41 |
|  | 30x | 0.15 | 0.26 | 0.03 | 0.06 | 0.12 | 0.20 |

P and S represent PhredEM and SeqEM, respectively. $G$ is the true genotype; the case $G = 2$ is omitted as it is barely seen for rare variants. MACs of 1, 5, 10, and 20 correspond to MAFs of 0.0005, 0.0025, 0.005, and 0.01, respectively, given the sample size of 1,000. All results are based on 5,000 replicates.

For common variants, we varied MAFs from 0.05 to 0.45 and assumed HWE in both data generation and model fitting. The results in Table II show that PhredEM outperformed SeqEM in the overall mis-called number as well as the stratified numbers. Overall, PhredEM correctly called 1–2 more genotypes at depth ≤ 10x and ~0.4 more at depth of 30x, with most of the improvement in major allele homozygotes which is the largest category. For both methods, the mis-called number declined substantially as the depth increased. The number mis-called increases as the MAF increases because the contribution to the likelihood (4) when *G*_*i*_ = 1 is independent of the *phred* score, which can easily be seen from (3). Furthermore, true minor allele homozygotes are more likely to be mis-called than major allele homozygotes due to the smaller prior probability of minor allele homozygotes.

**Table II:**
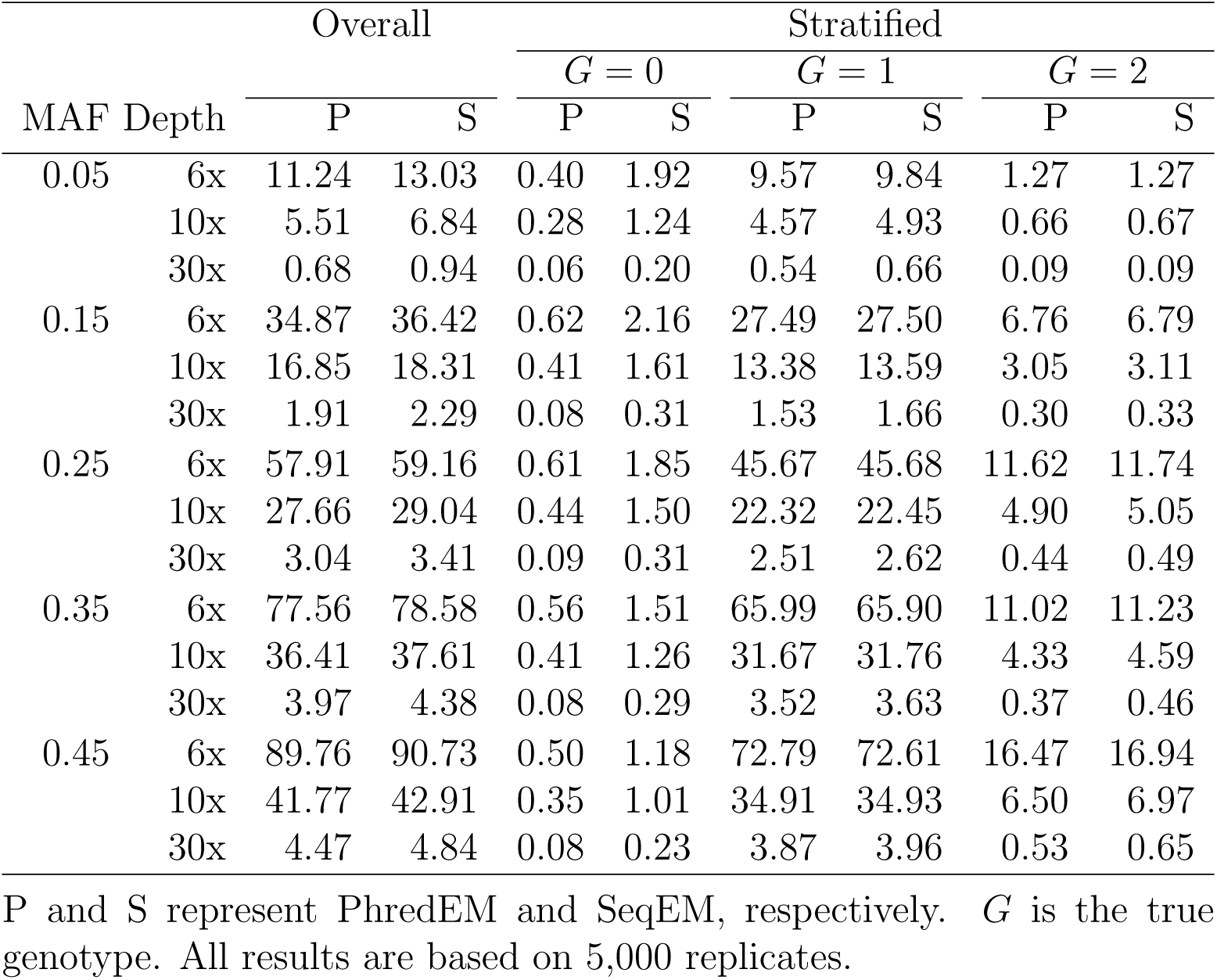
Number of mis-called genotypes for common variants in the simulation studies.

We further examined the *phred* scores at loci having genotypes that are called differently by PhredEM and SeqEM. In Table III, we displayed the average *phred* score associated with major and minor alleles at such loci, stratified by the true genotype (*G*) and genotypes called by PhredEM (*G*_P_) and SeqEM (*G*_S_). At loci with (*G*_P_, *G*_S_) = (0,1), regardless of the value of *G*, the major alleles tend to have high *phred* scores whereas the minor alleles tend to have low scores, explaining why PhredEM called these loci major allele homozygotes. The average *phred* scores for minor alleles are consistently lower under *G* = 0 than that under *G* = 1, because in the former case the minor alleles are all errors and in the latter case the minor alleles are a mixture of errors and true alleles. Similarly, for loci with (*G*_P_,*G*_S_) = (2,1), the major alleles tend to have low scores, which are lower under *G* = 2 than those under *G* = 1. In other cases when PhredEM called heterozygous genotypes, we observe high average *phred* scores for both major and minor alleles. These patterns of *phred* scores confirm that PhredEM worked as expected. While the results in Table III pertain to common variants, those for rare variants are similar and shown in Supplemental Table S3.

**Table III:** Average *phred* scores associated with called major (M) and minor (m) alleles at loci that are called differently by PhredEM and SeqEM in the simulation studies for common variants.

| MAF | Depth | $G = 0$ | | | | $G = 1$ | | | | | | | | $G = 2$ | | | |
| --- | --- | --- | --- | --- | --- | --- | --- | --- | --- | --- | --- | --- | --- | --- | --- | --- | --- |
|  |  | (0, 1) |  | (1, 0) |  | (0, 1) |  | (1, 0) |  | (1, 2) |  | (2, 1) |  | (1, 2) |  | (2, 1) |  |
|  |  | M | m | M | m | M | m | M | m | M | m | M | m | M | m | M | m |
| 0.05 | 6x | 37.1 | 9.0 | 37.2 | 36.0 | 37.1 | 12.7 | 37.2 | 38.7 | 38.5 | 35.8 | 33.1 | 37.3 | 38.5 | 34.2 | 26.0 | 37.4 |
|  | 10x | 37.1 | 9.1 | 37.0 | 36.3 | 37.1 | 12.8 | 37.1 | 38.7 | 39.0 | 37.2 | 22.6 | 38.2 | 38.4 | 33.1 | 13.8 | 36.7 |
|  | 30x | 36.9 | 9.4 | 37.0 | 36.2 | 37.0 | 15.1 | 37.3 | 38.6 | 37.8 | 36.4 | 26.4 | 36.9 | 38.7 | 34.0 | 9.9 | 37.2 |
| 0.15 | 6x | 37.1 | 8.7 | 37.0 | 35.9 | 37.0 | 11.9 | 37.1 | 38.8 | 38.5 | 36.4 | 8.7 | 37.4 | 36.8 | 33.2 | 7.5 | 37.1 |
|  | 10x | 37.1 | 8.8 | 37.1 | 36.0 | 37.1 | 11.7 | 37.1 | 38.7 | 38.6 | 37.0 | 9.7 | 36.9 | 35.2 | 37.0 | 8.5 | 37.2 |
|  | 30x | 37.1 | 9.1 | 37.1 | 36.5 | 37.0 | 11.9 | 37.1 | 38.7 | 38.7 | 37.0 | 10.4 | 37.5 | 36.5 | 36.4 | 8.2 | 37.2 |
| 0.25 | 6x | 37.1 | 8.6 | 37.5 | 36.2 | 37.2 | 11.3 | 37.1 | 39.0 | 38.9 | 36.2 | 9.7 | 37.1 | 38.2 | 32.1 | 8.2 | 37.2 |
|  | 10x | 37.2 | 8.8 | 37.1 | 36.1 | 37.1 | 11.0 | 37.1 | 38.8 | 38.6 | 36.9 | 10.3 | 37.4 | 36.9 | 36.1 | 8.4 | 37.0 |
|  | 30x | 37.1 | 8.8 | 37.2 | 37.0 | 37.0 | 12.3 | 37.0 | 38.6 | 38.2 | 37.1 | 11.8 | 37.4 | 36.8 | 37.3 | 9.3 | 37.1 |
| 0.35 | 6x | 37.1 | 8.5 | 36.6 | 37.5 | 37.1 | 11.0 | 37.1 | 39.0 | 38.6 | 33.9 | 25.6 | 37.1 | 38.5 | 28.2 | 18.3 | 37.2 |
|  | 10x | 37.1 | 8.6 | 37.1 | 35.3 | 37.2 | 10.7 | 37.1 | 38.7 | 38.5 | 36.0 | 22.0 | 37.2 | 38.4 | 31.9 | 13.2 | 37.1 |
|  | 30x | 37.1 | 9.2 | 37.3 | 35.9 | 37.1 | 11.4 | 37.1 | 38.7 | 38.3 | 36.8 | 14.7 | 37.4 | 36.8 | 35.6 | 9.1 | 37.0 |
| 0.45 | 6x | 37.1 | 8.3 | 31.7 | 38.0 | 37.2 | 11.1 | 36.6 | 38.9 | 39.0 | 37.0 | 10.8 | 37.1 | 37.3 | 35.9 | 8.4 | 37.2 |
|  | 10x | 37.1 | 8.6 | 36.6 | 37.1 | 37.3 | 11.1 | 37.1 | 38.5 | 38.9 | 37.0 | 10.4 | 37.1 | 36.5 | 37.0 | 8.5 | 37.1 |
|  | 30x | 37.1 | 8.8 | 37.1 | 36.6 | 36.9 | 11.8 | 37.2 | 38.6 | 38.3 | 37.1 | 9.7 | 37.4 | 37.3 | 37.7 | 9.0 | 37.1 |

$G$ is the true genotype. $(G_P, G_S) = (0, 1), (1, 0)$ , et al. represent loci that are called to be $G_P$ and $G_S$ by PhredEM and SeqEM, respectively. The results are based on 5,000 replicates.

### UK10K SCOOP DATA

We analyzed sequencing data from the Severe Childhood Onset Obesity Project (SCOOP) cohort, which was sequenced as part of the UK10K project. The sequenced SCOOP cohort consists of 784 UK Caucasian patients with severe, early onset obesity, and they were whole-exome sequenced by Illumina HiSeq 2000 with an average depth of ~60x. We first used SAMtools to generate pileup files from BAM files, filtering out reads that are PCR duplicates, with mapping score ≤ 30, or with improperly mapped mates. From the pileup files, we extracted read count data and *phred* scores. The distribution of the *phred* scores is shown in Figure 1(a).

Using the SCOOP sequencing data, we checked the fit of the logistic regression model in (2). First, we applied our screening algorithm to identify loci that were monomorphic (i.e., *π̂* = 0). At such loci, we could reliably treat all minor allele reads as errors. Assigning

*Y* = 1 and 0 for minor allele reads and major allele reads, respectively, we can determine the relationship between Pr(*Y* = 1) and the corresponding *phred* scores *Q*. To create a subset of such data that is computationally manageable, we randomly selected 1,000 monomorphic loci from each of the 22 chromosomes and randomly picked one individual from each locus, forming a dataset of 22,000 (*Y*, *Q*) pairs. Then, we fit the linear function of *phred* scores to ln{Pr(*Y* = 1)/Pr(*Y* = 0)} (i.e., the logistic regression model in [2]) and, as a gold standard, we also fit a smooth spline function of *phred* scores using the generalized additive model (GAM) [Wood, 2006]. Figure 1(b) shows the fitted curves and pointwise 95% confidence intervals from the two models. The two confidence regions overlap substantially for *phred* scores greater than 8, and the logistic regression fit fell within the 95% confidence region of the GAM for *phred* scores as low as 5. When *phred* scores were extremely low, the logistic regression model appeared to yield smaller base-calling error rates than the GAM, although it should be noted that only a very few *phred* scores this low were found in these data (see Figure 1a). Thus, we conclude that over the range of *phred* scores found in real data, the logistic model describes the relationship between *phred* scores and base-calling error rates well.

To facilitate the evaluation of PhredEM and especially the comparison with SeqEM, we first selected a set of genotypes that can serve as the gold standard. Specifically, we downloaded from the UK10K website the VCF files for the SCOOP cohort, which contained genotypes called by SAMtools and filtered by GATK. In addition, we excluded a variant if its average depth across samples is less than 20. We excluded a genotype whose genotype likelihood (on the *phred* scale) was ≤ 20 (i.e., genotyping error rate ≥ 0.01) and excluded a variant completely if it has more than 20% of genotypes with likelihood ≤ 20. These exclusion criteria ensured that all selected genotypes were called with particularly high quality. We thus refer to these genotypes as “true” genotypes. Since the loci with true genotypes were selected towards having high read depth, both PhredEM and SeqEM would perform well if applied to the original data. To create sequencing data with low or median depth, we adopted a subsampling scheme that sampled each read with equal probability. Finally, we applied PhredEM and SeqEM to call genotypes, assuming HWE for variants with MAF (calculated from true genotypes) ≤ 5% and allowing HWD for others. The computation time depends on the average depth. For example, it took approximately 30 hours and 128 MB memory on a single thread of an Intel Xeon X5650 machine with 2.67GHz for PhredEM to call the whole-exome genotypes in the 6x dataset.

The results of mis-called genotypes, averaged over all variants on chromosomes 1-22 and stratified by MAF ranges, are displayed in Table IV. For rare variants, the pattern in the number of mis-called genotypes by PhredEM and SeqEM agreed well with the results in the simulation section, with PhredEM generally producing more accurate genotype calls throughout the range of MAFs. The biggest difference occurred when the variants were relatively rare, i.e., MAF ∈ (0.001, 0.01]; when the average read depth was ~6x, PhredEM generated 1.8 more correct genotypes out of 758 individuals than SeqEM for loci with MAFs in this range. For more common variants, the differences between the two methods were smaller, possibly because *phred* scores at heterozygous loci are not informative; this also explains the increase in genotype-calling error rates with increasing MAF found throughout Table IV. The *phred* scores at loci with differently called genotypes by the two methods are summarized in Supplemental Table S4. These results exhibited the same patterns seen in the simulated data. In summary, all results show that PhredEM can improve the genotype-calling accuracy over SeqEM for real sequencing data in NGS studies.

**Table IV:** Number of mis-called genotypes in analysis of the UK10K SCOOP data (subsampled to achieve different depths).

| MAF | Depth | Overall |  |  | Stratified |  |  |  |  |  |  |  |  |
| --- | --- | --- | --- | --- | --- | --- | --- | --- | --- | --- | --- | --- | --- |
| | | $N$ | P | S | $G = 0$ | | | $G = 1$ | | | $G = 2$ | | |
| | | | | | $N_0$ | P | S | $N_1$ | P | S | $N_2$ | P | S |
| (0, 0.001] | 6x | 756.4 | 0.36 | 1.00 | 755.6 | 0.17 | 0.78 | 0.9 | 0.19 | 0.22 | 0 | 0 | 0 |
|  | 10x | 776.4 | 0.34 | 1.02 | 775.5 | 0.24 | 0.89 | 0.9 | 0.10 | 0.13 | 0 | 0 | 0 |
|  | 30x | 782.9 | 0.22 | 0.83 | 782.0 | 0.19 | 0.79 | 0.9 | 0.03 | 0.04 | 0 | 0 | 0 |
| (0.001, 0.01] | 6x | 757.7 | 1.99 | 3.80 | 753.1 | 0.57 | 2.30 | 4.5 | 1.33 | 1.41 | 0.1 | 0.09 | 0.09 |
|  | 10x | 776.1 | 1.78 | 3.43 | 771.4 | 0.50 | 2.06 | 4.6 | 1.19 | 1.28 | 0.1 | 0.09 | 0.09 |
|  | 30x | 782.3 | 1.52 | 2.23 | 777.6 | 0.37 | 1.05 | 4.6 | 1.09 | 1.12 | 0.1 | 0.06 | 0.06 |
| (0.01, 0.05] | 6x | 752.4 | 10.87 | 11.70 | 714.8 | 1.40 | 3.28 | 36.9 | 8.93 | 7.88 | 0.7 | 0.54 | 0.54 |
|  | 10x | 773.6 | 8.19 | 9.26 | 734.9 | 2.17 | 3.95 | 37.9 | 5.62 | 4.90 | 0.7 | 0.40 | 0.41 |
|  | 30x | 780.3 | 5.38 | 6.43 | 741.3 | 1.93 | 3.16 | 38.2 | 3.26 | 3.07 | 0.8 | 0.19 | 0.20 |
| (0.05, 0.1] | 6x | 750.5 | 20.10 | 20.34 | 647.1 | 1.31 | 1.85 | 98.9 | 16.34 | 15.94 | 4.5 | 2.45 | 2.55 |
|  | 10x | 773.4 | 11.73 | 12.15 | 666.9 | 1.23 | 1.65 | 101.8 | 9.00 | 8.95 | 4.6 | 1.50 | 1.55 |
|  | 30x | 781.0 | 2.28 | 2.42 | 673.6 | 0.59 | 0.63 | 102.7 | 1.32 | 1.40 | 4.7 | 0.37 | 0.39 |
| (0.1, 0.2] | 6x | 749.6 | 38.28 | 38.60 | 547.2 | 2.09 | 2.51 | 184.9 | 28.88 | 28.70 | 17.5 | 7.31 | 7.39 |
|  | 10x | 773.3 | 21.36 | 21.78 | 564.7 | 1.92 | 2.32 | 190.5 | 15.46 | 15.47 | 18.0 | 3.98 | 3.99 |
|  | 30x | 780.4 | 3.54 | 3.66 | 570.2 | 0.90 | 0.94 | 191.9 | 1.91 | 1.98 | 18.3 | 0.73 | 0.74 |
| (0.2, 0.3] | 6x | 748.3 | 62.59 | 62.88 | 424.6 | 2.76 | 3.09 | 276.2 | 46.23 | 46.10 | 47.5 | 13.60 | 13.69 |
|  | 10x | 772.7 | 33.93 | 34.32 | 438.5 | 2.66 | 2.96 | 285.2 | 24.68 | 24.72 | 49.1 | 6.59 | 6.64 |
|  | 30x | 780.3 | 4.80 | 4.92 | 443.0 | 1.15 | 1.22 | 287.6 | 2.60 | 2.63 | 49.7 | 1.05 | 1.07 |
| (0.3, 0.4] | 6x | 749.5 | 80.98 | 81.22 | 319.1 | 2.97 | 3.22 | 337.6 | 62.42 | 62.40 | 92.8 | 15.59 | 15.60 |
|  | 10x | 773.1 | 42.16 | 42.51 | 329.3 | 2.91 | 3.18 | 348.1 | 32.20 | 32.21 | 95.8 | 7.05 | 7.12 |
|  | 30x | 780.4 | 5.42 | 5.57 | 332.4 | 1.29 | 1.35 | 351.3 | 2.94 | 2.99 | 96.7 | 1.19 | 1.23 |
| (0.4, 0.5] | 6x | 748.6 | 96.08 | 96.21 | 229.5 | 4.39 | 4.60 | 368.7 | 75.70 | 75.60 | 150.3 | 15.99 | 16.01 |
|  | 10x | 773.3 | 49.95 | 50.08 | 237.0 | 3.34 | 3.50 | 380.8 | 39.45 | 39.30 | 155.5 | 7.16 | 7.28 |
|  | 30x | 780.6 | 6.99 | 7.12 | 239.5 | 1.34 | 1.39 | 383.9 | 4.38 | 4.41 | 157.2 | 1.27 | 1.32 |

$G$ is the true genotype. $N$ , $N_0$ , $N_1$ , and $N_2$ are the average numbers of individuals covered by at least one read. P and S represent PhredEM and SeqEM, respectively.

To gain more insights into the mechanisms of PhredEM and SeqEM, we listed in Table V the raw data at eight loci (from the subsampled dataset at 6x) that were called differently by PhredEM and SeqEM. Generally, base calls with low *phred* score are error-prone, and PhredEM treats these unreliable calls as likely errors when calling the genotype. By contrast, SeqEM depends heavily on the proportion of minor allele reads among the total reads and ignores the quality measure of each allele. For example, at Locus 1, the six major alleles were of high quality while the two minor alleles were likely to be errors. In this case, PhredEM distinguishes between alleles of different qualities and produced the correct genotype but SeqEM, which cannot account for low quality alleles, calls the incorrect genotype.

**Table V:**
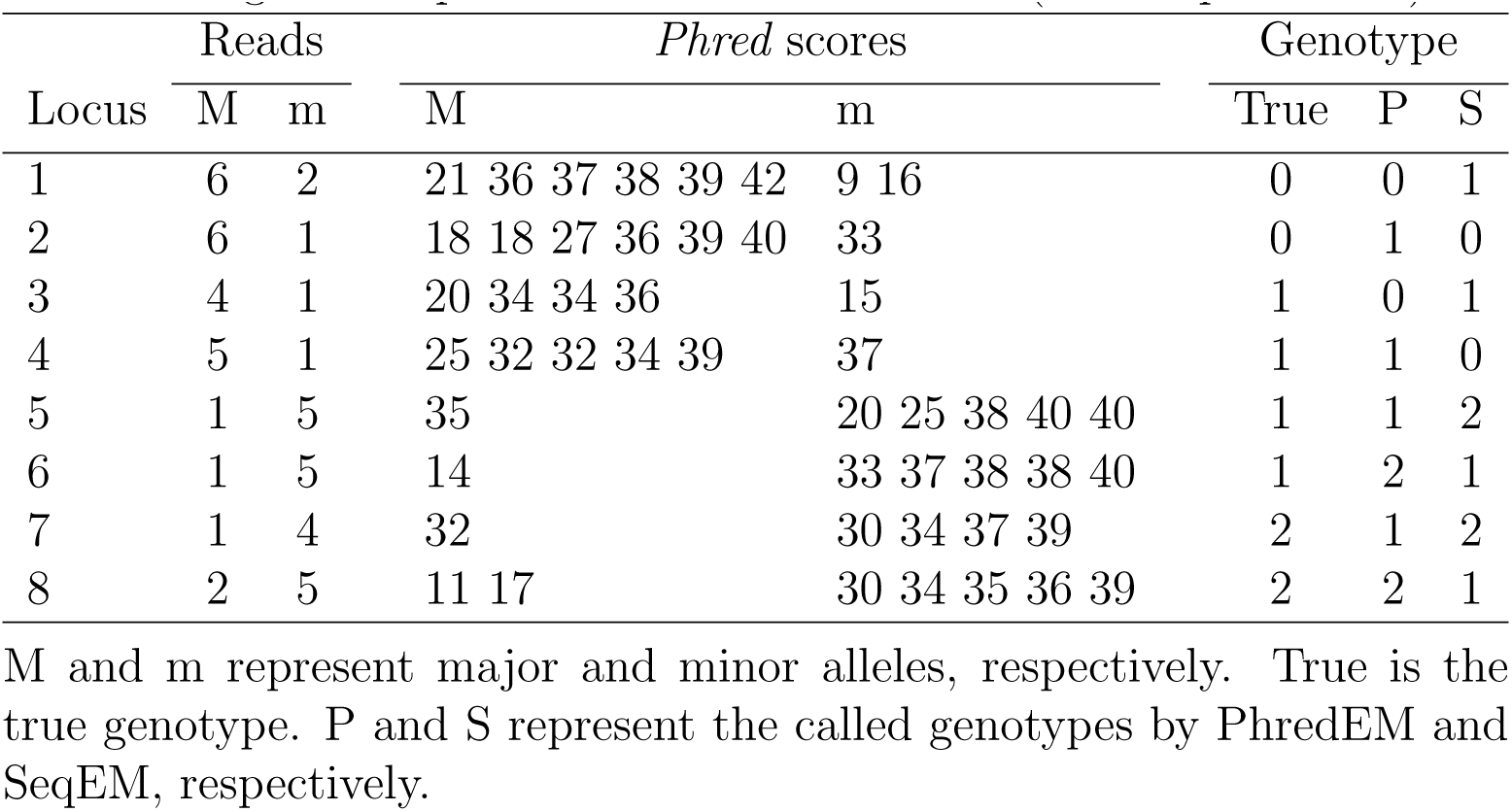
Eight example loci in the SCOOP data (subsampled to 6x).

## DISCUSSION

We have developed a *phred*-score-informed genotype-calling approach for NGS studies, called PhredEM. We also proposed a simple and computationally efficient screening algorithm to identify monomorphic loci. The PhredEM approach improves the accuracy of genotype-calling by estimating base-calling errors from both read data and *phred* scores, and by using all sequencing reads available without setting a *phred*-score-based quality threshold. PhredEM is closely related to the SeqEM approach, which can be viewed as a special case of PhredEM. We showed that the logistic model relating *phred* score to base-calling error rate used in PhredEM fits real sequencing data well.

In our logistic regression model (2), the *phred* score is the only predictor for the base-calling error. Other important predictors of base-calling quality could also be included. Because we estimate separate logistic regression parameters at each locus, covariates that are the same for each read (e.g., the particular nucleotides that constitute the minor and major alleles) are largely accounted for. One interesting covariate we have not considered is the position in the read [Brockman et al., 2008], although it is unclear whether this has an independent effect once the *phred* score is accounted for. We did not consider the mapping score as a possible covariate because there is not much variability in mapping scores [Li et al., 2008] (see Supplemental Figure S1). However, we recommend that PhredEM should be applied after excluding alignments with mapping scores less than 30.

We recommend using PhredEM with the HWD assumption as a default, because the model with HWD is more robust. After examining genotype frequencies obtained assuming HWD, a second pass of PhredEM could easily be made using the model assuming HWE. Our numerical studies (not shown) suggest that at medium or high read depth (≥10x), the estimated genotype frequencies based on the calls from PhredEM converged rapidly to their true values with increasing sample size even when assuming HWD.

We made some simplifying assumptions for PhredEM. First of all, the sample should consist of independent, unrelated individuals, which is essential to the likelihood in expression (4). A version of PhredEM could be constructed for trio data by modeling the joint genotypes of parents and offspring, for example, using the conditional-on-parental genotypes (CPG) approach of Schaid and Sommer [1993]. We also assume that base-calling errors are independent; in reality, the base-calling errors might be correlated due to factors such as library preparation and sequence context. We also assume that errors are symmetric, i.e. that the probability of a read for the major allele being mis-called as the minor allele is the same as the probability of the minor allele being mis-called as the major allele. Further, PhredEM assumes that all variants are biallelic. The biallelic assumption is reasonable because only a small fraction of SNPs have been verified to carry three or more alleles [Hodgkinson and Eyre-Walker, 2010]. In analyzing the SCOOP data, we deleted in advance all calls for bases that differed from the two most frequent bases at every locus. Finally, PhredEM makes no use of linkage disequilibrium information, and calls genotypes at each locus using only data from that particular locus. A haplotype-based version of PhredEM could easily be constructed, and may result in improved genotype-calling performance for common variants in very low-coverage data.

In summary, we developed PhredEM, an improved genotype caller which reduces the genotype-calling errors for NGS data. We also proposed a simple and computationally inexpensive algorithm for screening out loci that are estimated to be monomorphic. We showed that the proposed approach generates fewer incorrect calls than SeqEM regardless of the average read depth and sample size. Using the SCOOP data from the UK10K project, we demonstrated the capability of PhredEM to improve the genotype-calling accuracy in real sequencing data.

## APPENDIX

### EM ALGORITHM

In the EM algorithm, *G*_*i*_ (*i* = 1,…, *n*) is treated as missing. The complete-data log-likelihood has the form

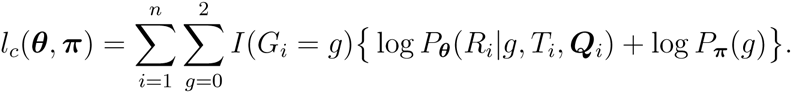

Let ***θ***^(*k*)^ and ***π***^(*k*)^ be the parameter values after the *k*th iteration. In the E-step of the (*k* + 1)th iteration, we evaluate *E*{*I*(*G*_*i*_ = *g*)|*R*_*i*_, *T*_*i*_, ***Q***_*i*_} for *g* = 0,1, 2, which can be shown to be

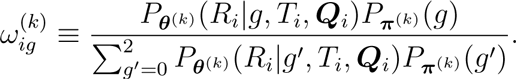

In the M-step, we maximize *l*_*c*_(***θ***, ***π***) with *I*(*G*_*i*_ = *g*) replaced by 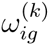. Specifically, under HWE we update *π* by a closed form 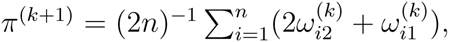, or under HWD we update *π* by the same *π*^(*k*+1)^ and update *f* by 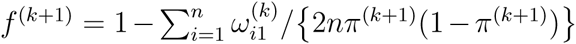. We use the one-step Newton-Raphson iteration to update ***θ***. We iterate between the E-step and M-step until the changes in the parameter estimates are negligible.

### PROOF FOR CONCAVITY OF *pl*(*π*)

First, we prove that, for fixed ***θ***, the function *h*(*π*) = log { ∑_*g*=0,1,2_ *P*_θ_(*R*|*g*,*T*, ***Q***)*P*_*π*_(*g*)} is concave. Under HWE, we write *h*(*π*) = log {*aπ*^2^ + *b*(1 — *π*)^2^ + 2c*π*(1 — *π*)}, where *a* = *P*_*θ*_(*R*|*G* = 2,*T*, ***Q***), *b* = *P*_***θ***_(*R*|*G* = 0,*T*, ***Q***), and *c* = (0.5)^*T*^. The second derivative of *h*(*π*) is

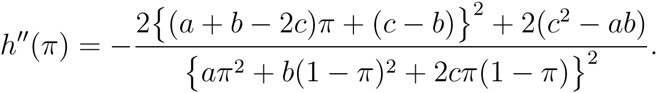

Because 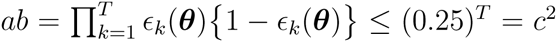,we obtainh“(*π*)≤ 0 and thus *h*(*π*)is a concave function of *π*.

Because the sum of concave functions is still concave, log *L*_*o*_(***θ***, *π*) is concave in *π* for fixed ***θ***. Because the pointwise supremum over ***θ*** preserves the concavity [Boyd and Vandenberghe, 2004], *pl*(*π*) is also concave.

### DISCLAIMER

The findings and conclusions in this report are those of the authors and do not necessarily represent the official position of the Centers for Disease Control and Prevention.

## ACKNOWLEDGMENTS

This study was supported by the University Research Committee (URC) Award at Emory.

